# Incorporating slow NMDA-type receptors with nonlinear voltage-dependent magnesium block in a next generation neural mass model: derivation and dynamics

**DOI:** 10.1101/2023.07.03.547465

**Authors:** Hiba Sheheitli, Viktor Jirsa

## Abstract

We derive a next generation neural mass model of a population of quadratic-integrate-and-fire neurons, with slow adaptation, and conductance-based AMPAR, GABAR and nonlinear NMDAR synapses. We show that the Lorentzian ansatz assumption can be satisfied by introducing a piece-wise polynomial approximation of the nonlinear voltage-dependent magnesium block of NMDAR current. We study the dynamics of the resulting system for two example cases of excitatory cortical neurons and inhibitory striatal neurons. Bifurcation diagrams are presented comparing the different dynamical regimes as compared to the case of linear NMDAR currents, along with sample comparison simulation time series demonstrating different possible oscillatory solutions. The omission of the nonlinearity of NMDAR currents results in a shift in the range (and possible disappearance) of the constant high firing rate regime, along with a modulation in the amplitude and frequency power spectrum of oscillations. Moreover, nonlinear NMDAR action is seen to be state-dependent and can have opposite effects depending on the type of neurons involved and the level of input firing rate received. The presented model can serve as a computationally efficient building block in whole brain network models for investigating the differential modulation of different types of synapses under neuromodulatory influence or receptor specific malfunction.

**Statements and Declarations:** The authors have no competing interests to declare.

**Funding:** We acknowledge support by H2020 Research and Innovation Action grants Human Brain Project SGA3 number 945539.

## Introduction

“N-methyl-D-aspartate (NMDA) and dopamine (DA) receptors and their interactions control an incredible variety of functions in the intact brain and, when abnormal, these interactions underlie and contribute to numerous disease states. These receptor interactions are relevant in such diverse functions as motor control, cognition and memory, neurodegenerative disorders, schizophrenia, and addiction.” (VanDongen, 2008)

Our work has branched out of an effort to develop a biophysical mechanistic framework for incorporating dopaminergic neuromodulation in whole brain network models (Sanz Leon et al., 2013), particularly with the aim to create personalized models for Parkinson’s disease (PD) patients, in the spirit of what has been done for the case of epilepsy (Jirsa et al., 2017). PD is associated with loss of dopamine in the dorsal Striatum due to degeneration of dopaminergic neurons of the Substantia Nigra Pars Compacta. The resulting dopamine depletion in the Basal ganglia (BG) is accompanied by changes in firing rates, as well as patterns of firing and levels of synchronization of neurons in the BG subcortical circuit (Bergman, 2021). However, the dynamic alterations are not merely a local phenomenon but are instead distributed across the macroscopic BG–thalamocortical networks (Tinkhauser et al., 2018); the increase in beta burst prevalence is accompanied by greater concurrence in time of beta burst activity across the network, during which highly stable phase locking is observed between the activity of cortical regions and BG nuclei (Yu et al., 2021). This motivates us to develop needed mathematical formulation for incorporating dopaminergic neuromodulatory processes in the Virtual Brain Modeling platform (Sanz Leon et al., 2013), in which parcellated brain areas are represented as a set of nodes on a network, each endowed by a dynamical system describing the aggregate neuronal activity in a given brain region. A first step towards this aim is to devise a way to capture the effects of dopamine on a mesoscopic (neuronal population) level.

We take as a starting point the seminal work in (Humphries et al., 2009) which argued that dopamine action on medium spiny neurons of the striatum can be captured in spiking neuron models as a scaling in the maximal conductances of NMDA, AMPA and GABA receptors. A similar framework was also suggested in (Durstewitz et al., 2000) for modeling dopamine action on pyramidal cells of the prefrontal cortex. More specifically, in the striatum, it is reported that dopamine enhances NMDA receptor (NMDAR) currents in neurons with D1-type dopamine receptors and attenuates AMPA receptor (AMPAR) currents in neurons with D2-type dopamine receptors, and that the balance between the two is at the core of the proper functioning of the striatum (Humphries et al., 2009; Lindahl & Kotaleski, 2016). Dopamine is also reported to enhance NMDAR currents via D1-receptors in the prefrontal cortex, while attenuating AMPAR currents and simultaneously enhancing GABA receptor (GABAR) currents (Durstewitz et al., 2000). As such, the effect of dopamine is, in fact, state-dependent and can switch from a predominantly net inhibitory effect (in low-activity states) to a net excitatory effect (in high-activity states) (Durstewitz et al., 2000). This special feature of dopamine action is facilitated by a unique property of NMDAR, which is its voltage-dependent nonlinear magnesium (Mg^2+^) block, a different mechanism than that governing the voltage-gated channels that generate the action potential. At the resting membrane potential, extracellular Mg^2+^ binds tightly to a site in the pore of the channel, preventing any ionic current flow, even in the presence of glutamate. It is only when the membrane is already depolarized enough (such as due to the opening of AMPAR channels) that the Mg^2+^ block is removed and ionic currents can flow (Jahr & Stevens, 1990; Nowak et al., 1984). As such, NMDARs are often referred to as “coincidence detectors”, allowing maximal current flow when two conditions are met: glutamate is present and the cell is already depolarized.

This Mg^2+^ block-induced nonlinear voltage-dependence of NMDAR conductivity, which is at the core of the role that NMDAR plays in neuronal dynamics, is often either omitted, for mathematical convenience, from traditional neural mass models (Wilson & Cowan, 1972) or reduced to a linearization around a local working point of mean membrane voltage (Brunel et al., 2001). More recently, a novel approach was put forward to derive exact mean field models of populations of quadratic-integrate-and-fire neurons by invoking the Lorentzian ansatz for describing the distribution of membrane voltage values (Montbrió et al., 2015). The resulting models, coined “next generation neural mass models”, offer the advantage of retaining the mean membrane voltage of the population as an explicit dependent variable, along side the mean firing rate, in the final mean field equations (Coombes, 2023). In the work we present here, we show that this allows us to incorporate in the derivation the full nonlinearity of the NMDAR conductance. This is done by rewriting the expression for the nonlinear NMDAR conductance as a piece-wise quadratic polynomial to maintain the applicability of the Lorentzian ansatz. This consequently permits the analysis of the effect of NMDAR across the full range of dynamical regimes of the system.

We consider a population of what is often referred to as an Izhikevich spiking neuron, which was derived as a canonical model for spiking neurons and, for different values of its four parameters, can reproduce different spiking and bursting behaviour of known types of neurons (Izhikevich, 2018). The Izhikevich neuron is basically a quadratic-integrate-and-fire (QIF) neuron with an added membrane recovery (adaptation) variable that provides negative feedback (Izhikevich, 2003). While the next generation neural mass models that exploited the Lorentzian ansatz where first derived for populations of simple QIF neurons (Byrne et al., 2017; Coombes & Byrne, 2019; Montbrió et al., 2015), the same approach was soon thereafter shown to be applicable for the case of Izhikevich neurons (Chen & Campbell, 2022) and the closely related QIF neurons with slow adaptation (Ferrara et al., 2023). We will here build on these efforts and extend this approach to include NMDAR type synaptic currents with nonlinear voltage-dependent conductance in the derivation of a mean-field model for a population of Izhikevich type neurons.

In the next section, we present the equations governing the population of neurons and the main steps involved in the derivation of the corresponding mean-field model. We then present the results of the numerical analysis of the dynamics for two example cases, that is, with parameter sets corresponding to two different types of neurons: 1) a population of excitatory regular spiking (cortical) neurons (Izhikevich, 2003) and 2) a population of inhibitory striatal neurons (Humphries et al., 2009). We generate bifurcation diagrams that show transitions in stability of constant firing rate states, as parameters vary, and emergence of oscillatory limit cycle solutions. We also present sample time series of simulations displaying the different types of possible oscillatory solutions. All results are contrasted to the case of omitting the nonlinear Mg^2+^ block from the NMDAR term, that is, the case of assuming linear NMDAR conductance-type current with the same slow synaptic decay time constant. The analysis shows that the NMDAR nonlinearity can lead to a voltage-dependent shift in the bifurcation diagram and a change in the relative size of the different operating regimes, with possible disappearance of the high constant firing rate states. In conjunction, in the oscillatory regime, the NMDAR nonlinearity can lead to alterations in oscillation amplitudes, frequency power profile, as well as the level of neuronal synchrony within the population. Ultimately, the presented mean-field model exhibits a rich range of dynamical behaviors and can serve as a building block for computationally efficient mechanistic whole brain models the explicitly incorporate the differential effects of neuromodulatory forces as a heterogeneous modulation of AMPAR, NMDAR and GABAR synaptic currents.

### Derivation of mean-field equations

We consider a population of all-to-all coupled Izhikevich neurons with conductance type synapses. The single neuron equations are of the form (Izhikevich, 2018):

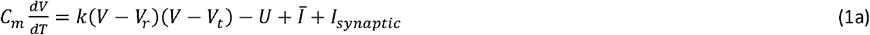

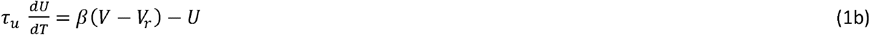

where *V* is the membrane potential, *U* is the recovery current, *C*_*m*_ is the membrane capacitance, *V*_*r*_ is the resting membrane potential, *V*_*t*_ is a threshold potential, *β* and *k* are scaling factors, *τ*_*u*_ is a time constant for the recovery variable, and *Ī* is an external applied current. The equations complemented with the following spike reset condition: *if V* > *Vpeak* : *V←V*_*reset*_ & *U←U + U*_*jump*_, whenever the membrane potential grows beyond the set peak value, *Vpeak it is reset back to the value, V*_*reset*,_ and the recovery current is augmented by a constant value *U*_*jump*_.

The synaptic currents can include excitatory AMPAR and NMDAR currents as well as inhibitory GABAR currents, such that: *I*_*synaptic =*_ *I*_*AMPA* +_ *I*_*NMDA +*_ *I*_*GABA*_

With *I*_*GABA=*_*G*_*G*_(*E*_*G −*_ *V*), *I*_*AMPA*_ = *G*_*A*_(*E*_*A −*_ *V*)

*& I*_*NMDA*_ = *G*_*N*_*B*(*V*)·(*E*_*N*_ − *V*) Where 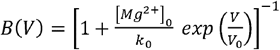

Here, *B*(*V*) is the nonlinear *Mg*^2+^ block, with [*Mg*^2+^]_0_=1*mM, k*_*0*_*= 3*.*57 &* 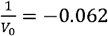, for normal operating conditions (Jahr & Stevens, 1990). (*E*_*G*,_ *G*_*G*_), (*E*_*A*,_ *G*_*A*_), (*E*_*N*,_ *G*_*N*_) correspond to the reversal potential and amplitude of conductance for GABAR, AMPAR and NMDAR, respectively. Ignoring the rise time for synaptic channel opening, the conductance amplitudes are governed by the following equations:

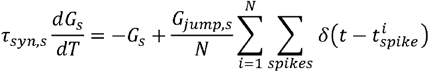

with subscripts *s* ∈{*A,N,G*}, referring to AMPAR, NMDAR and GABAR, respectively τ_*syn,s is*_ the synaptic decay time constant *G*_*jump,s*_ is the increment in conductance per spike received by a neuron, and *is* the total number of neurons in the population. After non-dimensionalization, the equations take the following simpler form:

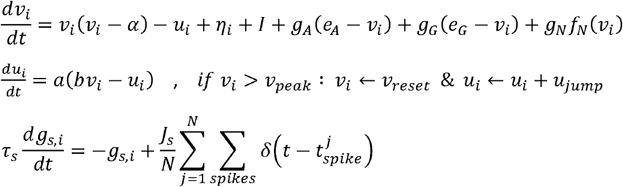

Where *i* ∈ *N* and 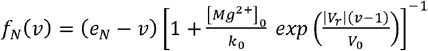

We have here included the *η*_*i*_ term, as a background current that introduces a heterogeneity among the neurons of the population. The scaled variables and parameters are related to the dimensional ones as follows:

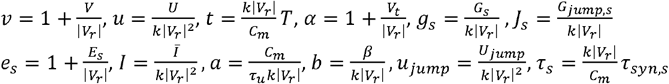

The expression for the *Mg*^2+^ voltage dependent nonlinear NMDAR current was, historically, obtained empirically by numerical fitting of experimental data (Jahr & Stevens, 1990; Nowak et al., 1984), so we can take a step back and instead rewrite it as a piece-wise polynomial. It can be seen in Figure 1 that *f*_*N*_ (*v*)can be well approximated as:

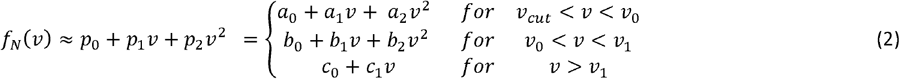

Given the reversal potential for NMDA is *E*_*N*_ = 0, *e*_*N*_ = 1, then the coefficients of the polynomial fit will only vary with the choice of *V*_*r*_. For the fit in figure 1, we used *V*_*r*_ = − 82.66*mV* (as in the first example case presented in the following Numerical Analysis section).

**Figure 1.**
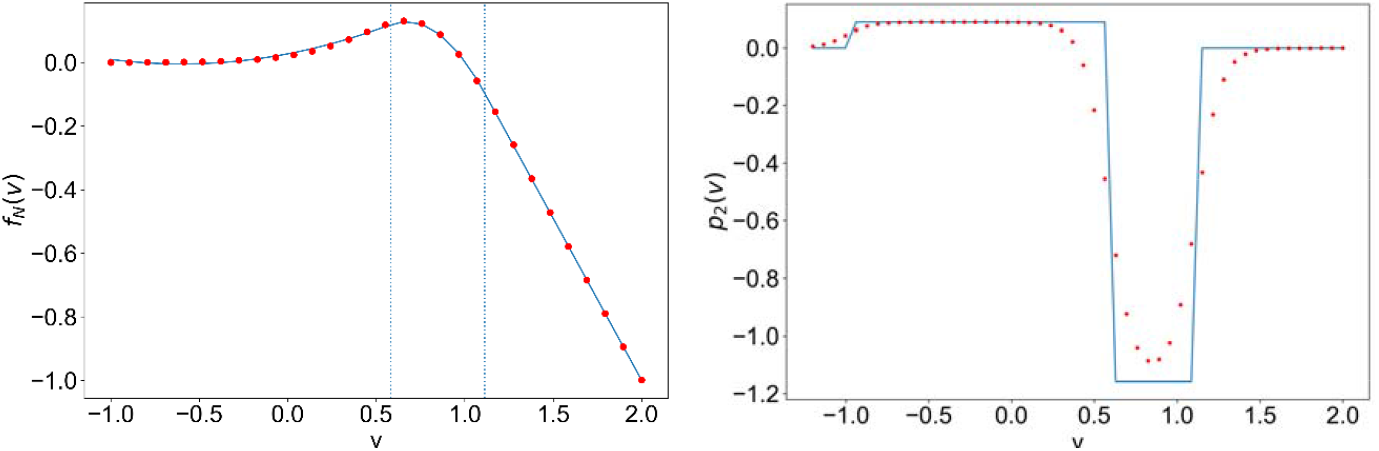
left: (··red) piece-wise approximation of (–blue) f(v). Right: (·· red) smooth approximation of (–blue) p_2_(v); for parameter values for the example case of excitatory regular spiking neurons reported in the next section.

The polynomial fit, as well as the discrete interval boundaries, were obtained by minimizing square errors over the range of values *v* ∈ [−1.2,2] while ensuring smoothness at the boundaries (that is, ensuring continuity in *f*_*N*_(*v*) and 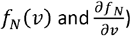)

Given that the time scale of the *u* dynamics is much slower than that of *v*, we can invoke the adiabatic approximation and consider that all neurons experience a common mean adaptation current, *u* = ⟨ *u*_*i*_ ⟩. We follow the same rigorous derivation presented in (Chen & Campbell, 2022), so for brevity, we only provide the main steps here and refer the reader to the latter reference for more elaborate details.

In the limit of *N*→∞, the distribution of the membrane potential is governed by the following continuity equation:

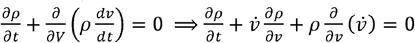

We assume a Lorentzian distribution for each of the membrane voltage and the heterogeneous background current:

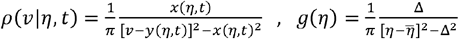

*x*(*η,t*) and Δcorrespond to the widths of the respective distribution, while *y*(*η,t*) and 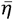 correspond to the respective centers.

The different terms of the continuity equation can be expanded as follows:

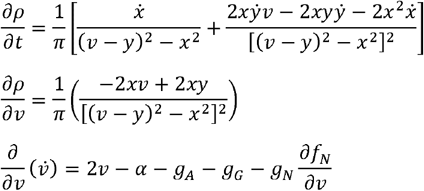

Plugging the above expressions in the continuity equation and balancing coefficients of *v*^*2*^&*v*, respectively, we get:

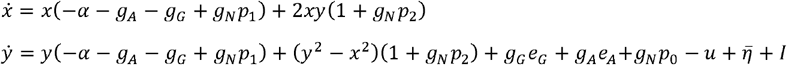

We then define *w* = *x* + *iy*, such that − *iw*^2^ = *i*(*y*^2^ − *x*^2^) + 2*xy* and 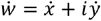, then the above 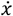 and 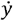 equations can be combined into:

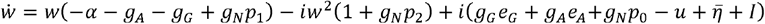

The firing rate can be obtained as the flux probability at *v*_*peak*_, as *v*_*peak*_ → ∞; for a given *η*:

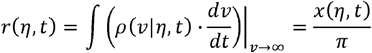

Here, we have used the fact that as *v* → ∞, *f*_*N*_(*v*) → *c*_0_ + *c*_1_*v*

Then the mean firing rate is: 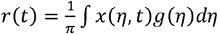

For a given value of *η*, the variable *y* can be computed as the Cauchy principal value of ∫ *ρ* | *η, tc*)*vdv*. Then, the mean membrane potential can be expressed as:

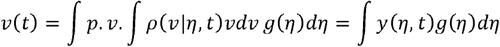

The residue theorem can be applied in a closed contour integral in the lower half complex plane containing the pole of 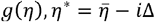, then: 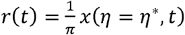 & *v*(*t*) = *y*(*η*= *η*\*, *t*), such that 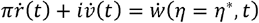.

Here, we have neglected the *ġ*_*N*_ contribution since the NMDAR dynamics is an order of magnitude slower than the dynamics.

Then, evaluating the equation for 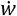 at 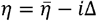, we arrive at the equations governing the mean firing rate and mean membrane potential:

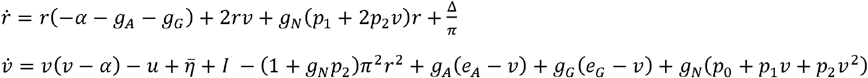

Noting that *p*_0_ + *p*_1_*v* + *p*_2_*v*^2^ = *f*_*N*_ (*v*) and 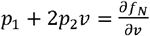, the final mean field equations can more compactly be written as:

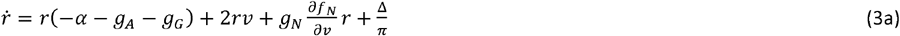

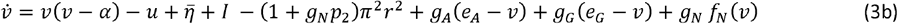

In addition, the equations governing the mean adaptation current and mean synaptic conductances take the following form:

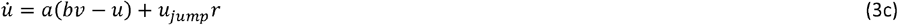

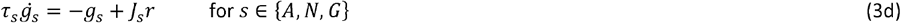

Here, the *r* in the *g*_*s*_ equation is the mean firing rate of the population in the case of recurrent synapses, or the mean firing rate received by the population in the case of an external input signal.

The coefficient *p*_2_ appearing on the right-hand side of the 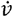 equation is defined piecewise as in eq.2, but to avoid discontinuity in the mean-field equations, we replace it with the following smooth approximate function:

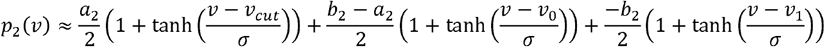

With *σ* = 0.15

The right panel of figure 1 shows the small error that we incur by making this step. We also replace the piece-wise expression for 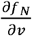 with its smooth equivalent by taking the derivative of the continuous *f*_*N*_ (*v*), such that:

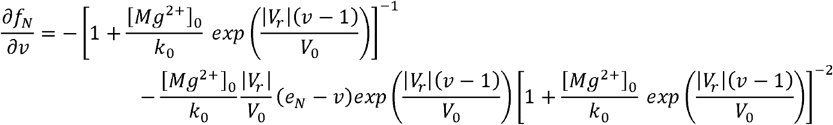

Finally, we note that there is a well-known link between QIF neurons and what is referred to as a *θ*-neuron which is described by a phase variable and can be used to describe excitable and spiking behavior. For a population of *θ*-neurons, the Kuramoto order parameter is defined as:

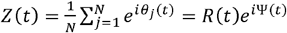

where *R* is a measure of the degree of synchrony within the network and Ψ is the average phase of the population, *R* = 0 & *R* = 1 correspond to a fully asynchronous and fully synchronized states, respectively (Byrne et al., 2017). A conformal transformation has been reported that allows one to move from the plane of the mean firing rate and mean membrane potential of a QIF population to the complex plane of the Kuramoto order parameter. Defining *W* = *πr* + *iv*, then the Kuramoto order parameter can be computed as: 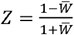

This allows us to have insight on the internal state of synchrony within the population and probe how it is affected with variation in the dynamical regime or parameter values, all the while staying on the level of a mesoscopic mathematical description of the population dynamics.

### Numerical Analysis

We numerically analyze the dynamics of the presented neural mass model for two example cases with typical sets of parameters. The bifurcation analysis as well as the numerical integration of the system of equations was performed using *PyDSTool* (Clewley et al., 2007), a time step of 0.1 was used for the timeseries simulation, and the power spectral density was obtained using the *scipy*.*signal*.*periodogram* python function.

### The case of excitatory regular spiking neurons

We analyze the resulting mean-field equations for the case of a population of excitatory regular spiking neurons with recurrent AMPAR synapses.

The equations for the Izhikevich regular spiking neuron are often written in the following form in the literature:

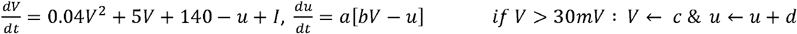

The part 0.04*V*^2^ + 5*V* + 140 came from fitting the spike initiation dynamics of a cortical neuron so that the membrane potential has *mV* scale, and the time has *ms* scale (Izhikevich, 2003). After factorization, this will correspond to the following parameter values for the original form of eq.1:

*C*_*m*_ = 1, *V*_*r*_ = − 82.656, *V*_*t*_ = − 42.344, *k* = 0.04, *a* = 0.02, *b* = 0.2, *u*_*jump*_ = 8 − *bV*_*r*_ =24.532, *I* = − *bV*_*r*_ = 16.532. For the corresponding non-dimensional system, this corresponds to *α* = 0.488. Note that here we shift the slow current variables to add a *V*_*r*_ term in the *u* equation for non-dimensionalization, such that we have a (*V* − *V*_*r*_) term to replace with *v* | *V*_*r*_ |. Accordingly, we add the non-zero external current term in the equation to correct for the shift in *u*.

We consider the case in which the population receives external excitatory input that drives both AMPAR and NMDAR synapses. The system of equations becomes: (from now on, we drop the bar from 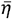)

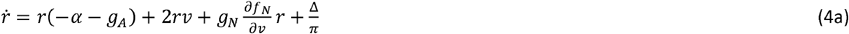

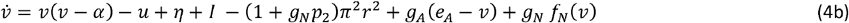

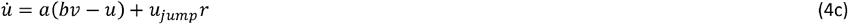

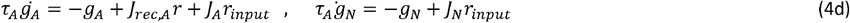

*J*_*rec*_,_*A*_ is the strength of the recurrent AMPAR synapses, whereas *J*_*A*_ & *J*_*N*_ are the strengths of the input AMPAR & NMDAR synapses and the corresponding reversal potentials are *E*_*A*_ = *E*_*N*_ = 0, such that *e*_*A*_ = *e*_*N*_ = 1. The synaptic time constants for AMPAR & NMDAR are taken to be 6*ms* & 160 *ms*, respectively, corresponding to *τ*_*A*_ 19.83 & *τ*_*N*_ 529.

For *V*_*r*_= −82.66, the coefficients for the *f*_*N*_(ν) piecewise fit are as follows: (we ν_*cut*_= −1.0) *a*_*0*_= 0.027, *a*_*1*_ = 0.106, *a*_*2*_ = 0.089, *b*_*0*_ =−0.396, *b*_*1*_= 1.559, *b*_*2*_ = 1.158, *c*_*0*_ = 1.038, *c*_*1*_ = −1.018, along with ν_*0*_= 0.582, ν_*1*_= 1.112.

For *r*_*input*_ = 0, we fix *J*_*rec,A*_ and analyze the stability of the equilibrium points as η is varied and obtain the bifurcation diagrams shown in figure 2. The plots show the value of the mean firing rate and mean membrane potential corresponding to the fixed-point solution of eqs.4a-d. For small enough η, the system has a stable equilibrium point as a low constant firing rate that loses stability through a Hopf bifurcation giving rise to a limit cycle solution of multiscale oscillations (of similar nature as that shown in figure 4). For intermediate values of η, the constant firing rate fixed point regains stability at high firing rate values, before losing stability again at a higher critical value of η and leading way to a stable limit cycle solution of the fast spiking-like oscillation type (as that shown in figure 5).

**Figure 2.**
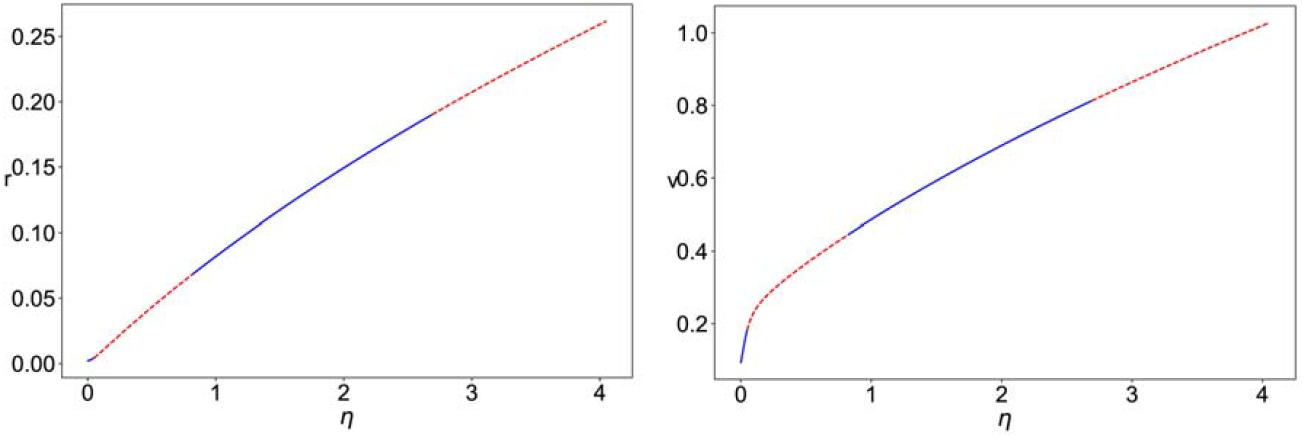
Bifurcation diagram for eqs.4a-d. with Δ= 0.002, J_rec,A_ = 6, r_input_ = 0. The plots show the value of mean firing rate (r, left) and mean membrane potential (v, right) corresponding to the stable fixed points (– blue) and unstable fixed point (--red) as a function of the value of the mean background current (η).

We then fix Δ, η, *J*_*rec,A*_, *J*_*A*_ & *J*_*N*_ and vary *r*_*input*_ Figure 3 shows the bifurcation diagrams for three configurations: *J*_*N*_ =0 (AMPAR only), *J*_*N*_ =3 with linear NMDAR (the nonlinear Mg^2+^ block is omitted), and *J*_*N*_ =3 with nonlinear NMDAR. In all cases, we have the following different dynamical regimes:

**Figure 3:**
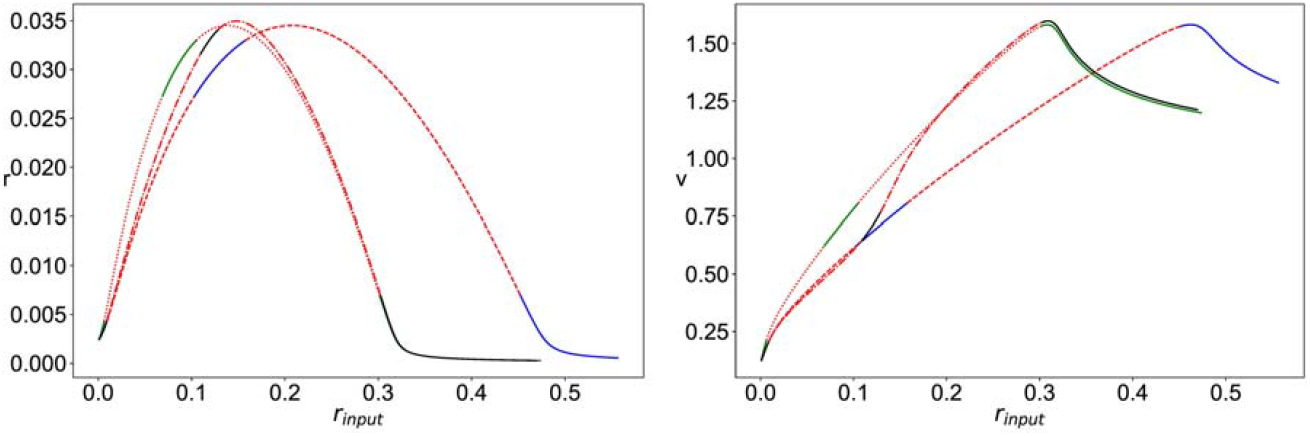
Bifurcation diagram for eqs.4a-d, with Δ= 0.002, η = 0.01, J_rec,A_= 6. The plots show the value of mean firing rate (r, left) and mean membrane potential (v, right) of the fixed points of the system as a function of the external input firing rate (r_input_), for three different cases: J_A_ = 6, J_N_ = 0 (stable –blue, unstable –red); J_A_ = 6, J_N_ = 3, with linear NMDA (stable – green, unstable ··red); J_A_=6,J_N_=3, with nonlinear NMDA (stable –black, unstable -.-red)

stable fixed point with a low constant firing rate, multiscale oscillation limit cycle, stable fixed point with high constant firing rate, and fast spiking-like oscillation limit cycle. With AMPAR only synapses, the latter regime persists for larger values of *r*_*input*_, that is, for the two cases of linear and nonlinear NMDAR, the curve is shifted to the left such that the fast spiking-like limit cycle loses stability at a significantly smaller value of *r*_*input*_. It can also be seen that in the presence of nonlinear NMDAR synapses, the regime of stable high firing rate fixed point is significantly reduced and shifted towards higher levels of *r*_*input*_ when compared to the case of linear NMDAR.

In figure 4, we show a sample simulation time series for the mean firing rate, the synchrony measure *R*_*sync*_, and the total synaptic current *I*_*synaptic*_ for sample parameter values in the regime of multiscale oscillations.

It can be seen that in the presence of nonlinear NMDAR, the oscillation is larger in amplitude with augmented slow and fast components. Figure 4 also shows the power spectrum density of the total synaptic currents indicating a downward shift in the main slow frequency and an upward shift in the dominant high frequencies.

**Figure 4.**
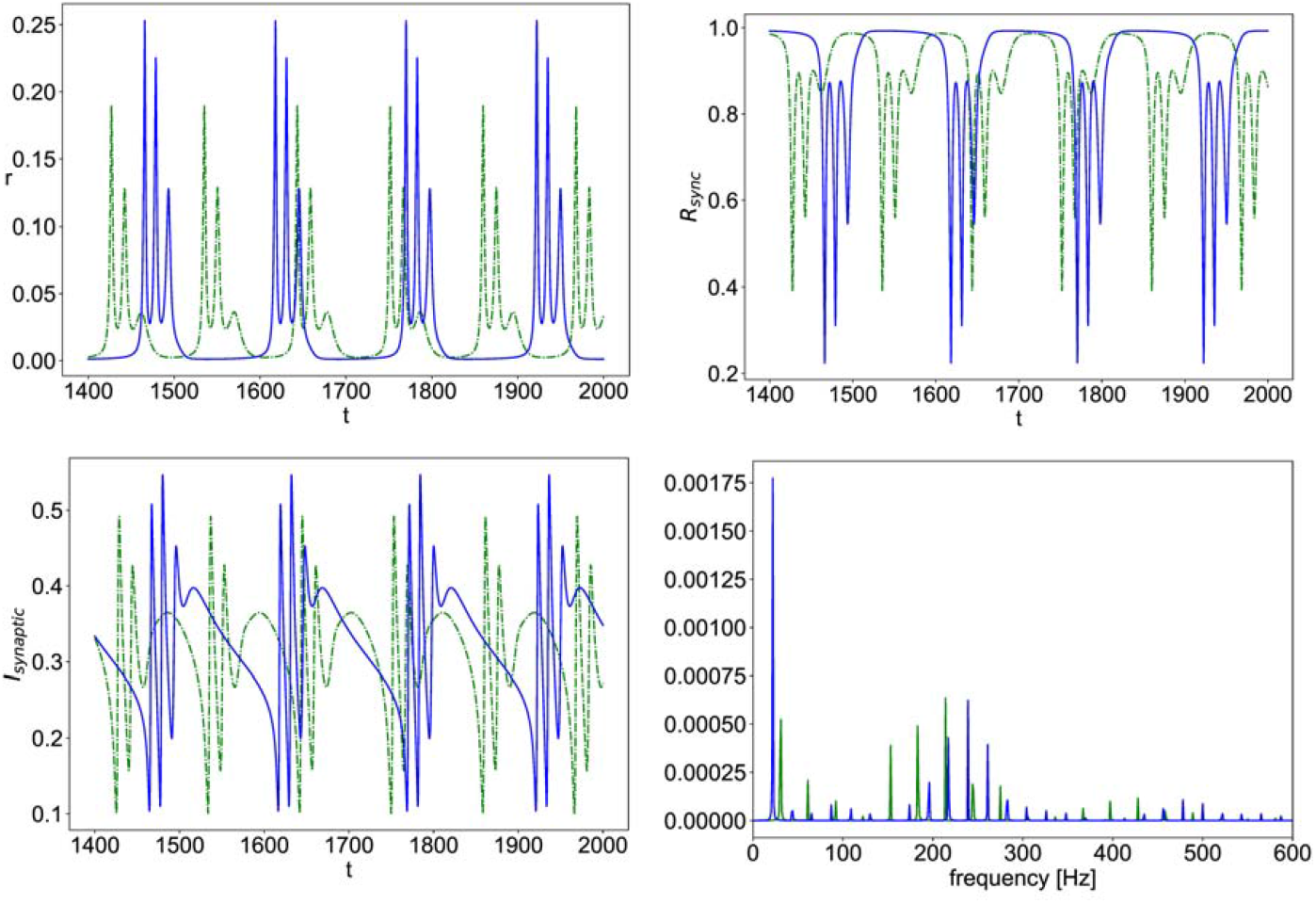
Simulated time series of eqs.4a-d with η = 0.01, Δ= 0.002, J_rec,A_ = 6, J_A_ = 6, J_N_ = 3, r_input_ = 0.06; the mean firing rate vs. time (r, top left), population synchrony measure vs. time (R_sync_, top right), total synaptic current vs. time (I_synaptic_, bottom left); power spectral density of the total synaptic current (bottom right); in all the plots, (–blue) is the case of nonlinear NMDA and (--green) is the case of linear NMDA.

**Figure 5.**
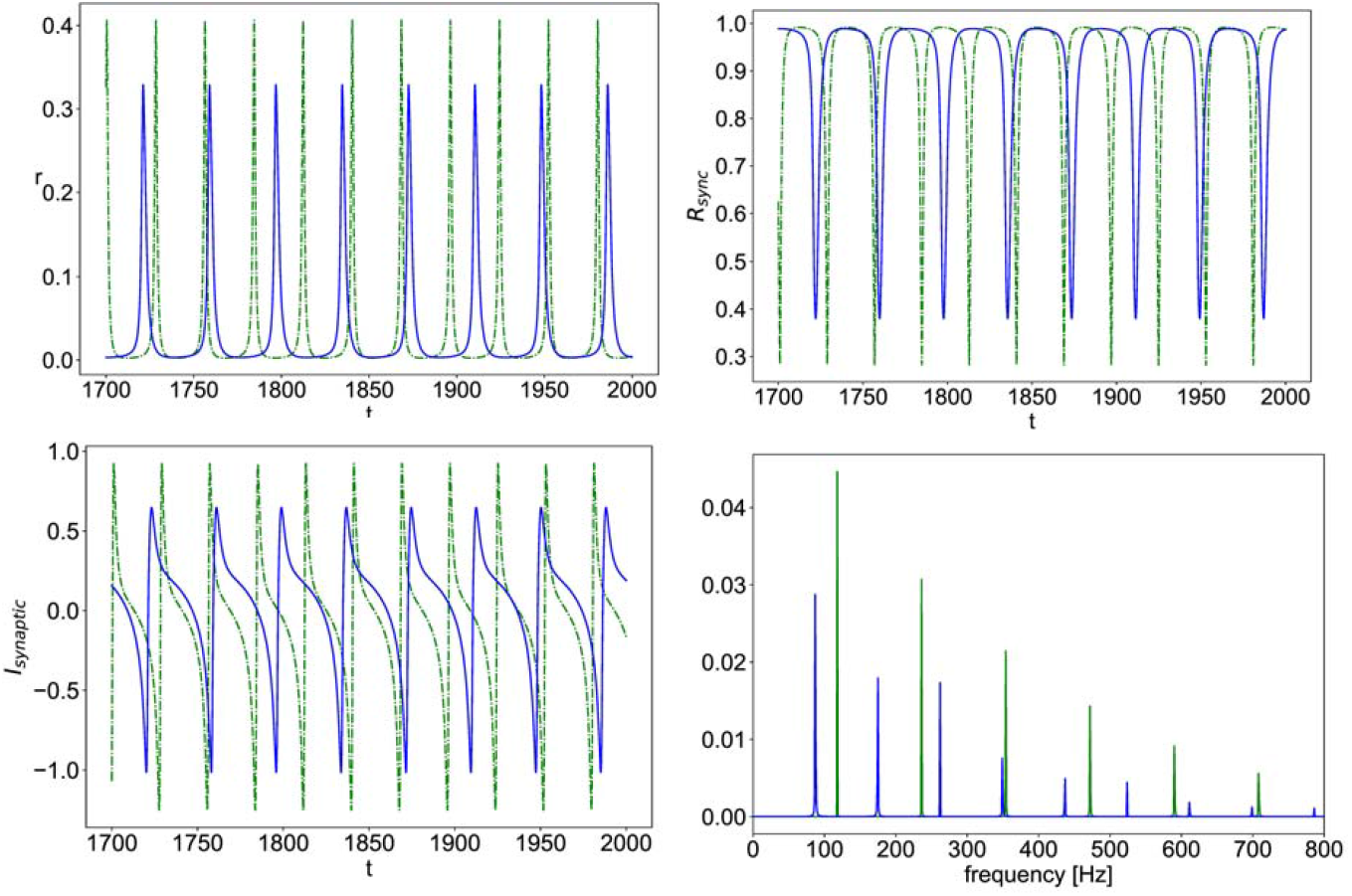
Simulated time of eqs.4a-d with η = 0.01, Δ = 0.002, J_rec_ = 6, J_A_ = 6, J_N_= 3, r_input_ = 0.17; the mean firing rate vs. time (r, top left), population synchrony measure vs. time (R_sync_, top right), total synaptic current vs. time (I_synaptic_, bottom left); power spectral density of the total synaptic current (bottom right); in all the plots, (–blue) is the case of nonlinear NMDA and (--green) is the case of linear NMDA.

Figure 5 shows another sample simulation data for parameter values in the regime of fast spiking-like oscillation. Here, the presence of nonlinear NMDA, instead, causes a smaller oscillation amplitude as compared to the case of linear NMDA, while still shifting frequency components of the total synaptic current towards lower values.

### The case of inhibitory striatal neurons

The second example case is that of a population of inhibitory striatal medium spiny neurons (MSNs), the specific parameters are taken from (Humphries et al., 2009) in which an Izhikevich neuron model was fit to capture neurocomputational properties of MSNs as well as the effect of dopamine on MSNs as a linear scaling in the AMPAR and NMDAR maximal conductances. The all-to-all coupled population of MSNs has recurrent inhibitory GABAR synapses and receives external excitatory AMPAR and NMDAR synapses. The resulting mean-field equations are:

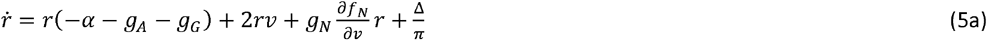

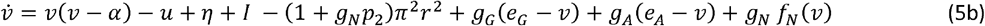

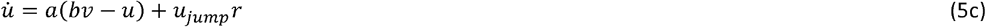

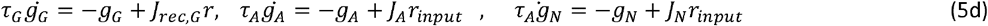

*J*_*rec,G*_, is the strength of recurrent GABAR synapses, whereas *J*_*A*_& *J*_*N*_ are the strengths of the input AMPAR & NMDAR synapses and the corresponding reversal potentials are *E*_*G*_ *= − 74 mV, E*_*A*_ *= E*_*N*_ *= 0*, such that *e*_*G*_*=0*.*075, e*_*N*_ *=1*. The synaptic time constants for AMPAR, NMDAR & GABAR are taken to be 6*ms*, 160*ms* &4*ms*, respectively, corresponding to τ_*A*_ *=*31.58, τ_*N*_*=* 842.1 & τ_*G*_*=* 21.05 after non-dimensionalization. The remaining parameters are as follows: *C*_*m*_ = 15.2, *V*_*r*_= −80, *V*_*t*_=−29.7, k =1, *a* = 0.01, *b* = −20, *u*_*jump*_= 91, *I* = 0.

For the corresponding non-dimensional system, we have: α = 0.629 and the coefficients for the *f*_*N*_*(ν)* piecewise fit are as follows: *a*_*0*_ = 0.0306, *a*_*1*_ = 0.113, *a*_*2*_ = 0.0914, *b*_*0*_ = −0.349, *b*_*1*_ = 1.464, *b*_*2*_ = −1.11, *c*_*0*_ = 1.039, *c*_*1*_ = −1.019, along with ν_*0*_ = 0.562, ν_*1*_= 1.118

Figure 6 shows the bifurcation diagram for the system of eqs.5a-d, with *r*_*input*_=0, as η is varied. The system has a stable constant low firing rate stable equilibrium that loses stability through a Hopf bifurcation for large enough η when fast spiking-like oscillations emerge.

**Figure 6.**
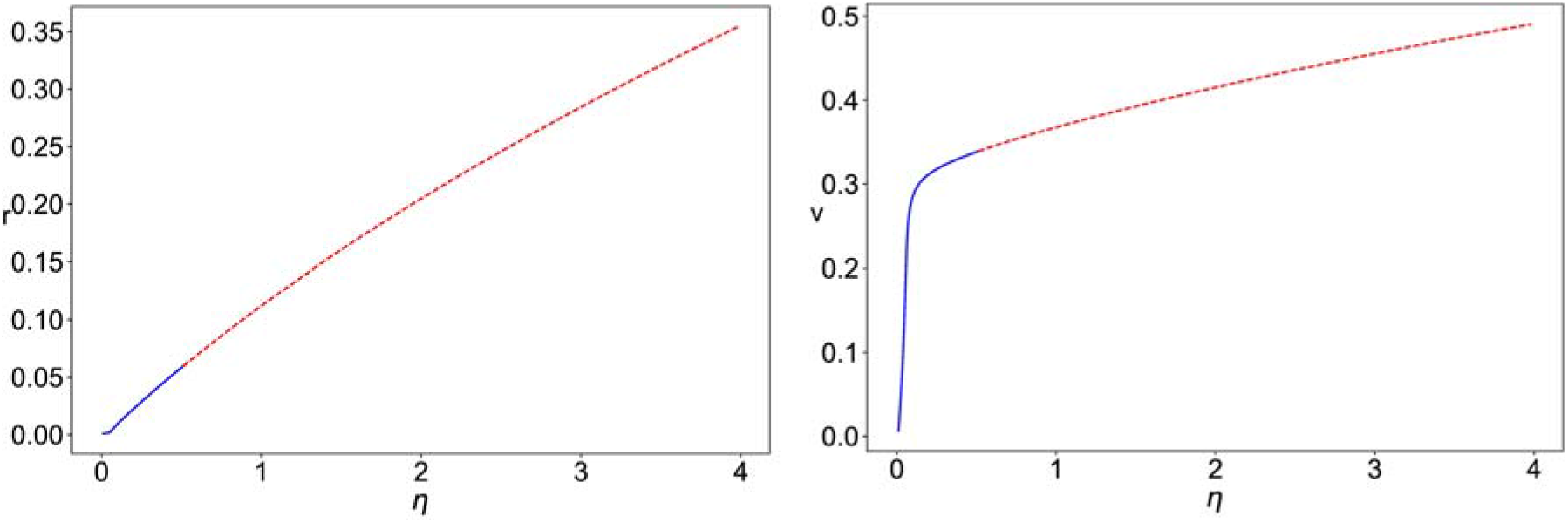
Bifurcation diagram for eqs.5a-d with Δ= 0.002, J_rec,G_= 1, r_input_ = 0The plots show the value of mean firing rate (r, left) and mean membrane potential (v, right) corresponding to the stable fixed points (– blue) and unstable fixed point (--red) as a function of the value of the mean background current (η).

In figure 7, we fix Δ, η, *J*_*rec,G*_, *J*_*A*_ & *J*_*N*_ and vary *r*_*input*_. It can be seen that the presence of nonlinear NMDAR allows for a regime of stable constant high firing rate solutions that do not exist for the case of linear NMDAR.

**Figure 7.**
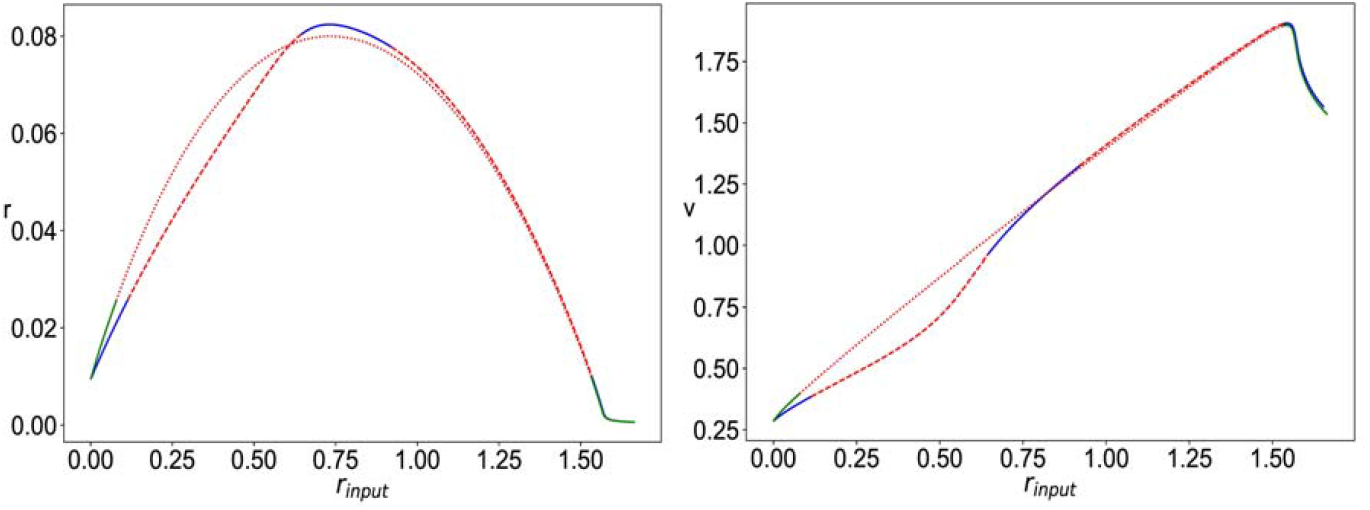
Bifurcation diagram for eqs. 5a-d with Δ= 0.002, η = 0.1, J_rec,G_= 1, J_A_ = 1.4, J_N_= 0.7; The plots show the value of mean firing rate (r, left) and mean membrane potential (v, right) of the fixed points of the system as a function of the external input firing rate (r_input_), for two different cases: with nonlinear NMDA (stable –blue, unstable –red) and with Linear NMDA (stable –green, unstable ··red)

Moreover, the sample simulation time series in figure 8 illustrates how nonlinear NMDAR leads to oscillations with a larger amplitude for *r*_*input*_ values that are below those of the high firing rate regime, along with a downward shift in the main frequencies of oscillation, as compared to the case of linear NMDAR. Whereas for *r*_*input*_ values that are above those of the high firing rate regime, nonlinear NMDAR has the opposite effect, leading to much smaller amplitude oscillations than those of the case of *r*_*input*_ value, along with a smaller downward shift in the frequencies, as shown in figure 9.

**Figure 8.**
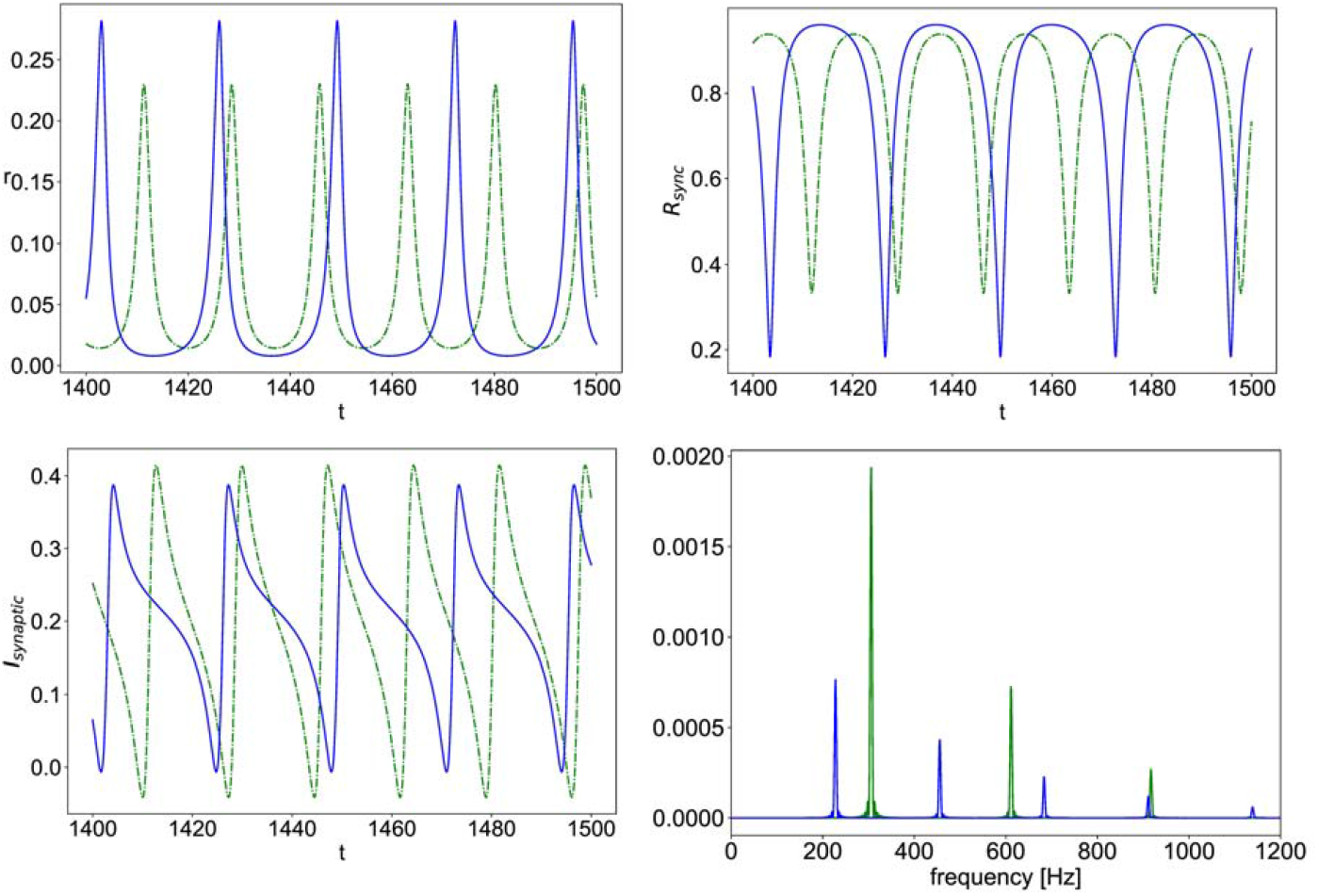
Simulation time series of eqs.5a-d with η = 0.1, Δ= 0.002, J_GABA_ = 1, J_AMPA_ = 1.4, J_NMPA_ = 0.7, r_input_ = 0.3; the mean firing rate vs. time (r, top left), population synchrony measure vs. time (R_sync_, top right), total synaptic current vs. time (I_synaptic_, bottom left); power spectral density of the total synaptic current (bottom right); in all the plots, (–blue) is the case of nonlinear NMDA and (--green) is the case of linear NMDA.

**Figure 9.**
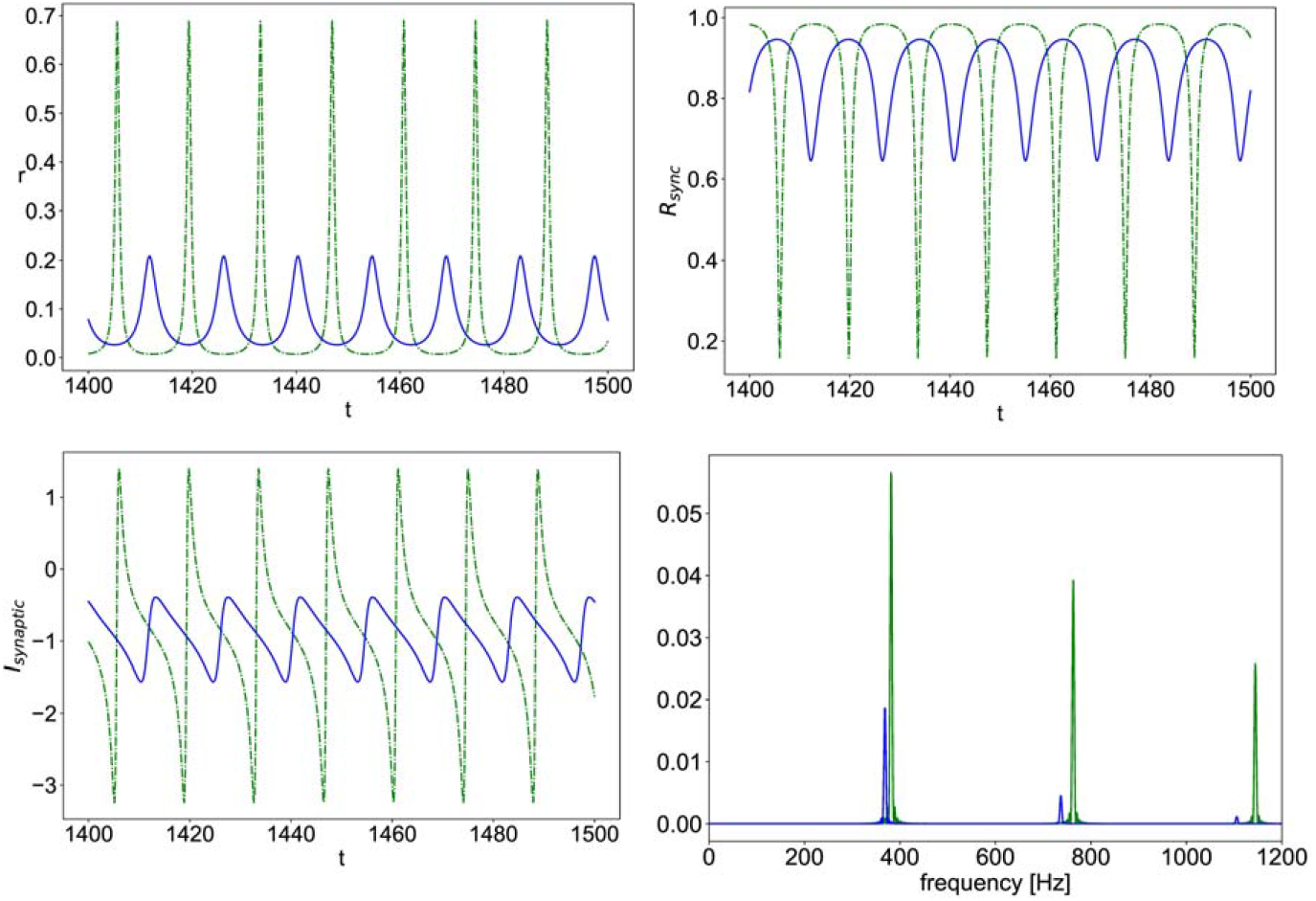
Simulation time series of eqs.5a-d with η=0.1, Δ=0.002,*J*_*GABA*_=1,*J*_*AMPA*_=1.4,*J*_*NMDA*_=0.7,r_*input*_=1.0; the mean firing rate vs. time (r, top left), population synchrony measure vs. time (*R*_*sync*_, top right), total synaptic current vs. time (I_synaptic_, bottom left); power spectral density of the total synaptic current (bottom right); in all the plots, (–blue) is the case of nonlinear NMDA and (--green) is the case of linear NMDA.

While an exhaustive parameter space exploration is beyond the scope of this work, it is worth noting that the effect of Nonlinear NMDA, as compared to the Linear case, was found to be qualitatively robust for a wide range of Δ & η values (results not shown here).

## Discussion

We presented a next generation neural mass model of Izhikevich-type neurons that includes NMDAR synaptic currents with nonlinear Mg^2+^ block, along with APMAR and GABAR conductance-based synapses. We have shown how rewriting the nonlinear Mg^2+^ block expression as a piece-wise polynomial maintains applicability of the Lorentzian ansatz in the mean-field model derivation. We have then performed numerical analysis of the resulting equations for two example cases. The first example involved a population of excitatory regular spiking neurons with recurrent AMPAR synapses and input AMPAR & NMDAR synapses. It was shown that the nonlinear NMDAR causes a shift in the constant high firing rate regime towards higher values of input and causes an amplification in amplitude of the multiscale oscillation solutions, while increasing the timescale separation by pushing the slow dominant frequencies downwards and the fast ones upwards. This is concurrent with larger intermittent dips in the population synchrony measure. Whereas, in the regime of fast spiking-like oscillations, nonlinear NMDAR exerts the opposite influence of decreasing the amplitude of oscillations in mean firing rate and those of the dips in population synchrony. The second example involved a population of inhibitory striatal medium spiny neurons with recurrent GABAR synapses and input AMPAR & NMDAR synapses. In this case, the nonlinearity in the NMDAR current gives rise to a constant high firing rate regime that is not present for the linear NMDAR case. This high firing rate regime separates two regimes of fast spiking-like oscillations, in which the effect of nonlinear NMDAR acts in opposite directions, for the lower input values, the amplitude of the fast oscillations is amplified due to the nonlinearity, while for the higher input values, the nonlinearity causes a significant decrease in the size of the oscillations and an associated curbing of the intermittent dip in population synchrony.

In summary, the nonlinearity of NMDAR synapses exerts a complex state-dependent effect on the population dynamics, that can manifest in opposite directions depending on neuronal type and level of input signal. This is in line with what is known on the complex state-dependent effect of dopamine action that utilizes the modulation of NMDAR maximal conductance as one of its routes of action. The presented model offers a computationally efficient building block for exploring the repercussions of complex NMDAR action, and associated neuromodulatory effects, on a mesoscopic and macroscopic brain level. By explicitly representing the different types of synaptic currents, not only with different timescales of action but also with the nonlinear state-dependent NMDAR feature, the model allows for the interrogation of mechanistic hypotheses on the role of imbalance in receptor or neuromodulator activity underlying mechanisms of brain disorders. That is, the formulation suggested here will permit going beyond the common simplifying assumptions of phenomenological sigmoidal input-output response functions, to a more biophysically grounded approach that expresses neuromodulatory action as a modulation of maximal conductance of the different synaptic currents involved. More specifically, the sigmoidal “firing rate vs. input current” response function, commonly used in classical mean-field models, is best suited to describe constant population firing rates in response to an input current that drives a near-constant deviation of membrane potential from its resting value (Abbott & Chance, 2005), a condition which is violated in the dynamical regime of oscillatory firing rate solutions representing complex burst-like activity as observed in the BG nuclei (Bergman, 2021). Moreover, the pharmacological sigmoidal function, recently proposed in the literature to account for neuromodulatory currents in whole brain network models (Joshi et al., 2017; Kringelbach et al., 2020), can only capture effects that enter as an additive current in target regions and falls short of capturing state-dependent parametric (i.e. multiplicative) effects as those reported in the case of dopaminergic action.

Our formulation allows for coupling multiple populations of different types of neurons, such as those of the BG nuclei, and opens the path to realistically investigate the effect of different dopamine levels as manifesting in a scaling of different maximal conductances to match known distribution of different receptor types (Lindahl & Kotaleski, 2016). Building the BG circuit using the proposed mean-field model will permit its embedding in the larger full brain network through connectome based whole brain models (Breakspear, 2017; Sanz Leon et al., 2013), and thus facilitate the study of emergent effects on a whole brain level, while preserving computational efficiency by avoiding spiking neurons simulation that would be required for detailed BG modeling (Humphries et al., 2006, 2018; Meier et al., 2022). The presence of a conformal mapping to go from mean firing rate and mean membrane potential to the Kuramoto order parameter variables offers the added advantage of probing alterations in average internal synchrony within regions while remaining on the level of mesoscopic description, this is particularly of relevance to disorders like PD in which the pathology manifests not only in changes in mean firing rate but also in that of coherence within the target neuronal populations (Bergman, 2021).

More broadly, NMDARs activity plays a central role in brain plasticity, learning and memory, the disruption of which is associated with impairment seen in a wide range of pathologies, such as Alzheimer’s disease, amyotrophic lateral sclerosis, Huntington’s disease, Parkinson’s disease, schizophrenia and major depressive disorder (Adell, 2020; Anticevic et al., 2012). We hope that the model proposed here will contribute to better representation and inclusion of nonlinear NMDAR action in the growing efforts on multiscale brain modeling, aiming to disentangle mechanisms of brain disorders, by bridging the gap between microscopic and macroscopic neuronal dynamics (D’Angelo & Jirsa, 2022; Deco & Kringelbach, 2014; Jancke et al., 2022; Joshi et al., 2017; Rolls et al., 2008; Shine et al., 2021)

